# Representations of fricatives in sub-cortical model responses: comparisons with human consonant perception

**DOI:** 10.1101/2022.10.24.513605

**Authors:** Yasmeen Hamza, Afagh Farhadi, Douglas M. Schwarz, Joyce M. McDonough, Laurel H. Carney

## Abstract

Fricatives are obstruent sound contrasts made by airflow constrictions in the vocal tract that produce turbulence across the constriction or at a site downstream from the constriction. Fricatives exhibit significant intra/inter-subject and contextual variability. Yet fricatives are perceived with high accuracy. The current study investigated modeled neural responses to fricatives in the auditory nerve (AN) and inferior colliculus (IC), with the hypothesis that response profiles across populations of neurons provide robust correlates to consonant perception. Stimuli were 270 intervocalic fricatives (10 speakers x 9 fricatives x 3 utterances). Computational model response profiles had characteristic frequencies that were log-spaced from 125 Hz to 8 or 20 kHz, to explore the impact of high-frequency responses. Confusion matrices generated by k-nearest-neighbor subspace classifiers were based on the profiles of average rates across characteristic frequencies as feature vectors. Model confusion matrices were compared them with published behavioral data. The modeled AN and IC neural responses provided better predictions of behavioral accuracy than the stimulus spectra, with IC showing better accuracy than AN. Behavioral fricative accuracy was explained by modeled neural response profiles, whereas confusions were only partially explained. Extended frequencies improved accuracy based on the model IC, corroborating the importance of extended high frequencies in speech perception.

## I. INTRODUCTION

### A. Fricatives

The broad goal of the current study was to investigate fricative sounds using a physiological model for responses of auditory-nerve (AN) fibers and midbrain neurons in the central nucleus of the inferior colliculus (IC) (Carney et al., 2015; Nelson and Carney, 2004; Zilany et al., 2014; Zilany et al., 2009). We used these computational models to test the hypothesis that response profiles across populations of neurons provide more robust correlates to consonant perception than the acoustic spectrum.

Fricatives are obstruent sound contrasts made by constricting airflow in the vocal tract, which produces turbulence across the constriction or a site downstream (Ladefoged, 1971; Ladefoged and Maddieson, 1996; Maddieson, 1984; 1991). The International Phonetic Alphabet (IPA) lists twelve places of articulation for fricatives, occurring at every place of articulation from labial to glottal. In the original UCLA Phonological Segment Inventory Database (UPSID) of the phonemic inventories of 317 languages dispersed across known language families, 296 have at least one fricative, with a mean of four fricatives (Maddieson, 1984; Maddieson, 1991). In a report on patterns in these inventories, Maddieson (1984) found that the coronal (dental through palatal) fricatives are among the most common phonemes occurring in phoneme inventories. While fricatives may exhibit voicing contrast sets (such as /s - z/, /ʃ - ʒ/), the distribution of voicing in fricatives is asymmetric. The most common fricative phonemic contrast is the dental or alveolar fricative, /s/, followed by /ʃ/ and /f/. The next most common are the voiced /z/, /ʒ/, and the voiceless velar fricative /x/. Paired sets of voiced-voiceless distinctions do not start to appear until a language has four fricatives, but, unlike other obstruent contrasts (plosives and affricates), voicing distinctions do not always occur in pairs. While the /z/ and /ʒ/ do not occur in the database without their voiceless counterparts, the voiced labial /β/, labiovelar /v/, interdental /ð/ and velar /ɣ/ appear with regularity without voiceless counterparts (Maddieson, 1984).

It has been long observed that the acoustic signature of any given fricative can vary significantly. Shadle (1985) has pointed out that fricatives may be best identified by formant transitions in the following vowels rather than by the fricative sound itself. In a cross-linguistic study of voiceless fricatives from seven languages, Gordon et al. (2002) found that spectral shape and center of gravity were important in distinguishing most fricatives, noting that when the fricatives had similar spectra and centers of gravity, the surrounding formant frequencies served to differentiate the fricatives. Fricatives also have been found to exhibit considerable intra/inter-subject variability (Jongman et al., 2000; Shadle, 1985; Shadle, 1990) as well as contextual variability (Narayanan et al., 1995). Temporal (duration and amplitude) and spectral properties have been used to identify fricative contrasts (Silbert and de Jong, 2008).

Much of the phonetic work on fricative sounds has been based on acoustics and aerodynamics, supported by speech perception studies (Chodroff and Wilson, 2020; Hughes and Halle, 1956; Jassem, 1979; McMurray and Jongman, 2011; Proctor et al., 2010; Shadle, 1985; Shadle et al., 1992; Strevens, 1960). Here, we asked how the neural representations of these sounds in the auditory pathway may address the gap between the variation found in fricative production and the accuracy of their identification.

Acoustically, fricatives, especially sibilant fricatives, are characterized by non-periodic energy at high frequencies. The most common ways used to quantify fricatives are the skewness, center of gravity, and dispersion of their acoustic spectra. Voicing affects the spectra of fricatives by introducing a low-frequency spectral peak and by reducing the amplitude of the higher-frequency noisy components (Crystal and House, 1988; Jesus and Shadle, 2002; Klatt, 1976). Place of articulation also affects the spectral properties of fricatives and is subject to individual and contextual variability that does not impede fricative perception (Chodroff and Wilson, 2020; 2022; Hughes and Halle, 1956; Jesus and Shadle, 2002; Jongman et al., 2000; Shadle, 1990; Shadle et al., 1992). While intensity is difficult to define or measure in speech, an early study of intensity by Stevens (1960) as muscular effort and air pressure demonstrated that there is little correlation between intensity and high-frequency energy in fricatives. Ladefoged and Maddieson (1996) noted that, in comparison to other sounds, significant demands are involved in the articulation of fricative sounds, in terms of the precision needed to shape the particular airstream channel and in maintaining the target shape over the duration of the sound.

Sibilant (/s/, /z/, /ʃ/, /ʒ/) and non-sibilant (all other) fricatives differ in spectra, amplitudes, and durations (Behrens and Blumstein, 1988; Evers et al., 1998; Hughes and Halle, 1956; Shadle, 1990; Strevens, 1960). Sibilant fricatives involve a noise source that is downstream of constriction, thus their spectrum is shaped by the resonant frequencies of the cavity anterior to the constriction (Stevens, 1998), whereas non-sibilant fricatives channel turbulence only at the site of constriction (Catford, 1977). This difference results in sibilant-fricatives typically exhibiting a midfrequency spectral peak, the result of the intense high-frequency turbulence caused by the airstream hitting the teeth, as opposed to non-sibilant fricatives that have a flat spectrum (Jongman et al., 2000; Shadle, 1990; Shadle et al., 1992). Sibilant fricatives have relatively higher amplitudes than non-sibilants.

The spectral differences described above have been demonstrated using statistical techniques such as spectral moments (Forrest et al., 1988; Jesus and Shadle, 2002; Jongman et al., 2000); however, such temporal and spectral features are not independent and are commonly variable across subjects and contexts (Chodroff and Wilson, 2022; Jongman et al., 2000; Narayanan et al., 1995; Shadle, 1990). For example, it is common that phonologically voiced fricatives undergo substantial devoicing in certain contexts (Haggard, 1978; Stevens et al., 1992), and phrase-final segments tend to be lengthened (Klatt, 1976). Despite this and other variability, listeners have been shown to be accurate in the perception of these contrasts (Cutler et al., 2004; Gallun and Souza, 2008; Pisoni and Luce, 1986; Woods et al., 2010).

While much phonetic work has focused on the articulation and acoustic analysis of fricatives, minimal investigation of their representation in the auditory system has been done (McMurray and Jongman, 2011). The broad goal of the current study is to investigate the modeled neural responses of fricatives from several speakers and repeated utterances (intra and inter-subject variability) in the auditory-nerve (AN) and midbrain (central nucleus of inferior colliculus, IC) responses, and compare them to results of a published behavioral task (Gallun, and Souza, 2008). The aim is to understand the perceptual salience of fricatives despite their variability in production. The hypothesis is that average-discharge-rate response profiles across populations of neurons provide more robust correlates to consonant perception than does the acoustic spectrum.

### B. Auditory-nerve and inferior colliculus modeling

The relationship between acoustics and neural encoding is complex due to strong non-linearities in the auditory periphery. Computational models for auditory neurons provide a strategy to describe these nonlinear transformations (Osses Vecchi et al., 2022). AN responses are affected by basilar membrane compression (Rhode, 1971), synchrony capture (Deng and Geisler, 1987; Miller et al., 1997; Young and Sachs, 1979), synaptic adaptation (Goutman and Glowatzki, 2007; Moser and Beutner, 2000; Raman et al., 1994; Westerman and Smith, 1984), and rate saturation (Sachs and Abbas, 1974; Yates, 1990; Yates et al., 1990). A computational model that includes these nonlinearities (Zilany et al., 2014; Zilany et al., 2009) was used in this study to characterize the responses of the population of AN fibers to fricatives.

The profile of discharge rates across the population of AN fibers is not a simple representation of the stimulus spectrum due to saturation of average rates. This limitation of average-rate representations is often addressed by studying phase-locking of AN fibers to temporal fine structure (e.g., Young and Sachs, 1979). However, not only is the temporal fine structure of fricative stimuli highly complex, the importance of high-frequency components in these stimuli limits the utility of phase-locking to temporal fine structure for encoding the full frequency spectrum of these sounds. Furthermore, the representation of the temporal fine structure of higher-frequency components of speech sounds is not likely to be critical in exciting auditory neurons at higher levels of the auditory pathway, such as the midbrain. However, relatively low-frequency fluctuations in AN responses, which are strongly elicited by noisy sounds such as fricatives, are important in exciting and suppressing neurons at higher levels of the auditory pathway (for a review refer to Carney, 2018). The depth of the low-frequency fluctuations in AN response varies along the frequency axis: near spectral peaks, the fluctuations are relatively shallow, whereas in spectral valleys, the fluctuations are deepest. The sensitivity of most neurons in the auditory midbrain to low-frequency fluctuations thus provides a potential neural representation for the spectrum of complex sounds. The robust transformation of important spectral *features*, such as peaks, slopes, and valleys of the spectrum, into AN fluctuation cues, and ultimately the average discharge rates of fluctuation-sensitive midbrain neurons, motivated the hypothesis that midbrain response profiles may provide a more robust representation of fricatives than that based directly on the values of spectral magnitudes.

It is interesting to consider the representations of complex sounds at the level of the midbrain, a nearly obligatory synapse for ascending projections from several brainstem nuclei (Schreiner and Winer, 2005). In the current study, the AN model (Zilany et al., 2014) provided the input to models for two types of midbrain neurons. Frequency tuning in the IC is inherited from the tuning of neural inputs and is described by a characteristic frequency (CF, the frequency eliciting a response at the lowest sound level). Additionally, most IC neurons are tuned for amplitude-modulation frequency, as described by modulation transfer functions (plots of average discharge rate as a function of sinusoidal amplitude modulation frequency) and a best modulation frequency (the modulation frequency that elicits the greatest excitation or suppression in the modulation transfer function) (Krishna and Semple, 2000; Joris et al., 2004; Nelson and Carney, 2007). Most IC neurons have best modulation frequencies in the range of voice pitch (Langner, 1992).

The majority of IC neurons are either excited (band-enhanced, BE) or suppressed (band-suppressed, BS) by amplitude modulations across a band of modulation frequencies spanning the best modulation frequency (Kim et al., 2020; Kim et al., 2015). This study explored representations of fricatives in the average rates of populations of model BE and BS neurons (Carney et al., 2015; Nelson and Carney, 2004). These two cell types respond differently to the profile of low-frequency fluctuations that are set up in AN responses. BE neurons are excited by low-frequency fluctuations of their inputs, thus these neurons respond most strongly when they are tuned to frequencies near spectral slopes, which elicit large low-frequency amplitude fluctuations in the responses of narrowband peripheral filters and AN fibers. In contrast, BS neurons are excited by signals with minimal fluctuations, thus they respond most strongly when they are tuned near spectral peaks, which elicit relatively small fluctuations in AN responses due to capture of inner-hair-cell responses by harmonics near spectral peaks (Carney, 2018). The response rates of both types of IC neuron are also affected by spectral levels near their characteristic frequencies or at low-frequencies, which drive auditory neurons through the “tails” of tuning curves. Because of the differences in the response properties of BE and BS IC neurons, we compared the ability to classify fricatives based on the population responses of each type. And because these two types of neurons can be considered as “opponent” cell types (Kim et al., 2020), we also explored classifier performance based on the combined responses of BE and BS cell types.

The current study tested the proposed hypothesis that behavioral accuracy in a fricative-identification task involving stimuli with and without spectro-temporal degradation (Gallun and Souza (2008) could be better explained by neural representations of the sounds at the levels of the AN and IC than by the spectra of the acoustical signals. We tested the hypothesis by computing average-rate responses of model AN fibers and IC neurons to fricatives, and then estimating performance in a categorization task based on these model response profiles, as compared to that based on the spectral energy profile. Categorization performance based on either the neural models or the acoustic spectrum was also compared to performance of listeners in Gallun and Souza (2008).

### C. Extended High-Frequency Hearing

Hearing loss is conventionally defined as elevated thresholds within the frequency range of 125 Hz to 8 kHz (WHO, 2008). The limitation to 8 kHz in conventional pure-tone audiometry is based on the finding that much of the studied phonetic information is provided by frequencies below 6 kHz (Vitela et al., 2015). However, there is growing evidence suggesting that acoustic information in the higher-frequency regions affects speech intelligibility, particularly in noisy environments (Badri et al., 2011; Hunter et al., 2020; Levy et al., 2015; Monson et al., 2019; Moore et al., 2017; Polspoel et al., 2022; Zadeh et al., 2019). Hearing loss at the extended high frequencies is common with aging and noise exposure and may reflect cochlear synaptopathy, which occurs first at higher frequencies (Liberman et al., 2016). In the current study, we compared the accuracy of identifications based on the acoustical signal, on model-AN responses and on model-IC neurons, using information limited to 8 kHz vs. an extended frequency of 20 kHz. As previously stated, much of the phonetic information that has been studied is limited to 6 kHz, therefore we hypothesized that accuracies of identifications based on an acoustic analysis would not differ with the limitation of frequencies. On the other hand, because high-frequency auditory neurons respond across a wide range of frequencies, in part due to the low-frequency tails of AN tuning curves (Kiang and Moxon, 1974), limiting the computational models to 8 kHz was expected to reduce accuracies of the identifications based on responses of model AN fibers and IC neurons, in line with the aforementioned evidence of the importance of high-frequency hearing in speech intelligibility.

## II. METHODS

### A. Stimuli

The recordings were done at a sampling frequency of 44.1 kHz, with 16-bit resolution, using a Marantz PDM661 Shure cardioid lavalier microphone, reported to have a frequency response that is flat within +/- 5 dB from 50 Hz to 17 kHz. The stimuli were the four sets of English fricatives, contrasting in place of articulation and voicing: labio-dental (/f/, /v/), interdental (/θ/, /ð/), alveolar (/s/, /z/), palatal-alveolar (/ʃ/, /ʒ/) plus the glottal fricative (/h/) in an intervocalic aCa (vowel-fricative-vowel) context. Ten native English speakers (5 males and 5 females) were recorded in a quiet room while seated comfortably in the presence of the experimenter with the microphone clipped approximately 10 cm below the mouth. The nine stimuli were read off a prepared list; participants were instructed to repeat each item three times, thus twenty-seven utterances were collected from each speaker.

The 270 utterances (10 speakers x 9 fricatives x 3 utterances) were segmented on Praat (Boersma and Weenink, 1992-2022) and demarcation of the onset, offset and midpoint of the initial vowel and fricative was determined for each utterance by visual inspection of the spectrogram and waveform. Formant frequencies and voicing cues, when possible, were used as guides.

Each of the aCa tokens was then scaled so that the level of the initial /a/ mid-section (60 ms) was 65 dB SPL. This scaling allowed the preservation of the relative levels of sibilant vs. non-sibilant fricatives. In the current study, model responses were obtained for entire utterances, and then model responses were analyzed over the time course of the mid-fricative segment (see Table 1 for mid-fricative sound levels).

**Table 1.** Mid-fricative sound pressure level (SPL) in dB; values are mean, standard deviation (SD), and range of root-mean-square (RMS). Levels shown were based on the mean of the 30 utterances (10 speakers x 3 repetitions) of the 60-ms mid-fricative sections.

| Mid-Fricative | RMS Mean (SD), dB SPL | RMS Range, dB SPL |
| --- | --- | --- |
| /s/ | 50.94 (5.03) | 39.40-63.47 |
| /z/ | 53.89 (4.91) | 47.74-63.96 |
| /ʃ/ | 51.71 (5.26) | 42.82-64.55 |
| /ʒ/ | 55.18 (5.04) | 46.36-66.12 |
| /f/ | 37.83 (4.18) | 28.39-45.09 |
| /v/ | 54.39 (5.58) | 42.89-64.18 |
| /θ/ | 37.84 (4.81) | 29.92-47.11 |
| /ð/ | 53.27 (5.77) | 39.00-64.47 |
| /h/ | 48.62 (5.31) | 38.61-56.64 |

For comparison of acoustic analysis to model neural responses, spectral information for each of the 270 utterances was computed for the 60-ms duration mid-fricative segment (extracted with a rectangular window) using a discrete-time Fourier transform to compute spectral magnitudes at individual frequencies that matched the log-spaced CFs used in the neural-response models. This log frequency spacing was similar to others used in speech analyses, e.g., Mel or Bark frequency axes, but the strategy used here allowed an exact match of the frequency channels between the acoustical and neural model representations.

Guided by Gallun and Souza (2008), four spectrally-degraded, speech-test conditions were also created: fricatives were processed by digital filtering with one-, two-, four-, or eight-channels that spanned from 176 to 7168 Hz. Intermediate cutoff frequencies for the channels were logarithmically spaced. The filtered segments were then degraded by randomly multiplying each sample of the filter outputs by +1 or −1, before summing the signals together to create the processed stimuli (Schroeder, 1968). This processing obscured the spectro-temporal information to varying degrees, with the one-channel signal providing the least spectral and temporal information, and the two-, four- and eight-channel signals providing increasing amounts of spectro-temporal information from the fricative sounds.

### B. Modeling

The average rates of neurons in response to each of the 270 aCa utterances were simulated using models for the AN and IC (Carney et al., 2015; Nelson and Carney, 2004; Zilany et al., 2014). The model response profiles, to the original and spectrally degraded stimuli, were computed for 60-ms duration segments centered within each fricative. The full model population had 50 CFs, log-spaced from 125 Hz to 20 kHz. In response to each utterance, the average discharge rate over the 60-ms duration for each of the 50 CFs for a given set of model neurons formed a feature vector that was used to classify the fricative stimuli (classifier details are below). There are nine possible classes corresponding to English fricatives (/f/, /v/, /θ/, /ð/, /s/, /z/, /ʃ/, /ʒ/, /h/). To understand the importance of extended high frequencies (beyond 8 kHz), we compared classifier performance for the unprocessed condition based on all 50 channels of the 20 kHz model to that based on the subset of 41 channels that extended from 125 Hz to 8 kHz. The average discharge rates of AN and IC models across the 41 or 50 CF channels were used as feature vectors for the classifier described below. Comparison of the confusion matrices of the spectra vs. neural-model responses in the extended (up to 20 kHz) vs. the conventional (8 kHz) frequency conditions highlighted the potential contribution of the high-frequency channels. Additionally, classification performance for processed stimuli, i.e., with spectro-temporal degradation, was compared to that for the unprocessed stimuli. For all of the processed stimulus conditions, the analysis and models had 50 CFs from 125 Hz to 20 kHz.

The AN model used in the current study was for low-threshold high-spontaneous rate fibers, which are the majority of AN fibers (Liberman, 1978). The AN model takes into consideration key non-linearities, previously mentioned, including compression, rate saturation, adaptation, and synchrony capture (Zilany et al., 2014). The original model was based on the physiological responses of the AN in cat; here, we used a version with a middle-ear filter and sharper peripheral tuning to represent the human ear (Ibrahim and Bruce, 2010), based on physiological and psychophysical measures (Shera et al., 2002).

The same-frequency inhibition-excitation (SFIE) model (Carney et al., 2015; Nelson and Carney, 2004) was used to model the responses of BE and BS IC neurons. The AN model provided the input for both BE and BS IC model neurons in the form of time-varying rate functions, convolved with functions representing excitatory or inhibitory postsynaptic responses, which differed for the two cell types (Carney et al., 2015). The postsynaptic potential time constants and delays were set to produce BE responses with a best modulation frequency of 100 Hz (Carney and McDonough, 2019). This best modulation frequency is near the center of the distribution of IC best modulation frequencies (Kim et al., 2020; Krishna and Semple, 2000; Nelson and Carney, 2007). Note that the modulation tuning of model and IC neurons is relatively broad (Q ≈ 0.5-1, Nelson & Carney, 2007). The BS model receives an inhibitory input from the output of the BE model. The code used for the simulations is available at https://osf.io/6bsnt/.

### C. Classifier Analysis

A k-nearest-neighbors classifier-based analysis was used to generate confusion matrices, from which the accuracy in identifying each fricative and the confusions between fricative contrasts were computed. Confusion matrices were constructed based on feature vectors that included the 50 (or 41) CF channels of spectral magnitudes or response profiles for AN, BE, BS, or combined BE + BS model neurons. For the combined BE + BS model, 25 CFs that were log-spaced from 125 Hz to 20 kHz were used for each model cell type, to create a single feature vector with 50 entries, matching the length and frequency range of those for the other models. Confusion matrices were then compared to published behavioral data for fricatives derived from the consonant-identification task based on the stimuli described above (Gallun and Souza, 2008).

To avoid overfitting of the classifier, the whole dataset (270 spectral/neural population responses) was divided using a cross-validation technique (20 folds) into training and testing subsets. The choice of 20 folds was based on the dataset size, and to reduce bias and variance in the models (Kohavi, 1995). In this case, the dataset was divided into 20 subsets, 19 were used to train the classifier and one was used to test the classifier. This process was repeated 20 times, to use all possible training and testing combinations, and the average accuracy (percent-correct prediction) was calculated. Training the model was based on the feature vector of each of the data points in the training dataset, with a label identifying the class (fricative) of data. When the classifier was trained, the testing dataset was used to estimate the classes based on the training data.

Additionally, the classifier model was tested for overfitting by randomly splitting the dataset into training and test sets with size of 230 and 40 samples respectively. The classifier was trained using the training dataset and tested with both training and testing datasets for three trials. For all trials and for all type of responses, the difference between mean absolute error for training and testing sets was negligible (less than 2%).

The classifier type was a subspace, ensemble classifier that uses k-nearest-neighbor learners. In the current study, the number of learners was 30 (spectral/neural responses per fricative), and the subspace dimension was 50 (or 41 in an 8 kHz model), corresponding to the length of the feature vector. The MATLAB’s Statistics and Machine Learning Toolbox was the platform used for the classifier analysis.

For the unprocessed conditions, training and testing were done using the unprocessed condition dataset, whereas for the processed conditions, training was done for the unprocessed condition and testing was done using the processed condition. The aim behind this strategy was to mimic the process of the behavioral response in normal-hearing individuals, for which degraded speech was presumably compared to auditory signatures in memory to reach a decision. This strategy was supported by the improvement of identification of unintelligible degraded speech and the enhancement in neuronal population responses after exposure to intact sublexical cues (Al-Zubaidi et al., 2022).

The behavioral data was derived from the phoneme-recognition task by Gallun and Souza (2008). For each condition, the task included 64 tokens consisting of 16 aCa syllables (/b, d, g, p, t, k, f, θ, s, ʃ, v, ð, z, ʒ, m, n/) spoken by four speakers presented at 65 dB SPL. Ten normal-hearing young participants participated by selecting the correct syllable in a 16-syllable choice forced paradigm displayed on a touch screen.

## III. RESULTS

Population average-rate responses to the unprocessed mid-fricative stimuli are shown as a function of model CF for sibilants (Fig. 1) and non-sibilants (Fig. 2). Additionally, the variation in the spectra and response profiles across the different processing conditions of a selected voiced sibilant (/asa/), a voiceless sibilant (/aʒa/), and a voiceless non-sibilant (/afa/) fricative, are illustrated in Figs. 3-5. The rate profile across AN fibers can be interpreted as a relatively straightforward representation of the acoustic magnitude spectrum, with peaks and valleys in AN rates aligned with corresponding features in the spectrum, but with an overall rate profile that is shaped by the middle-ear filter to accentuate mid-frequency responses. However, the model IC response profiles differ markedly from the acoustic spectra. Model BE neurons have the highest rates near spectral slopes, where AN responses have relatively large fluctuations. Model BS neurons are excited by the channels with the least fluctuation, near the peaks in the spectra, but are suppressed by fluctuations associated with spectral slopes. Thus, the BS rate profile is “sharper” than the AN rate profile.

**Figure 1.**
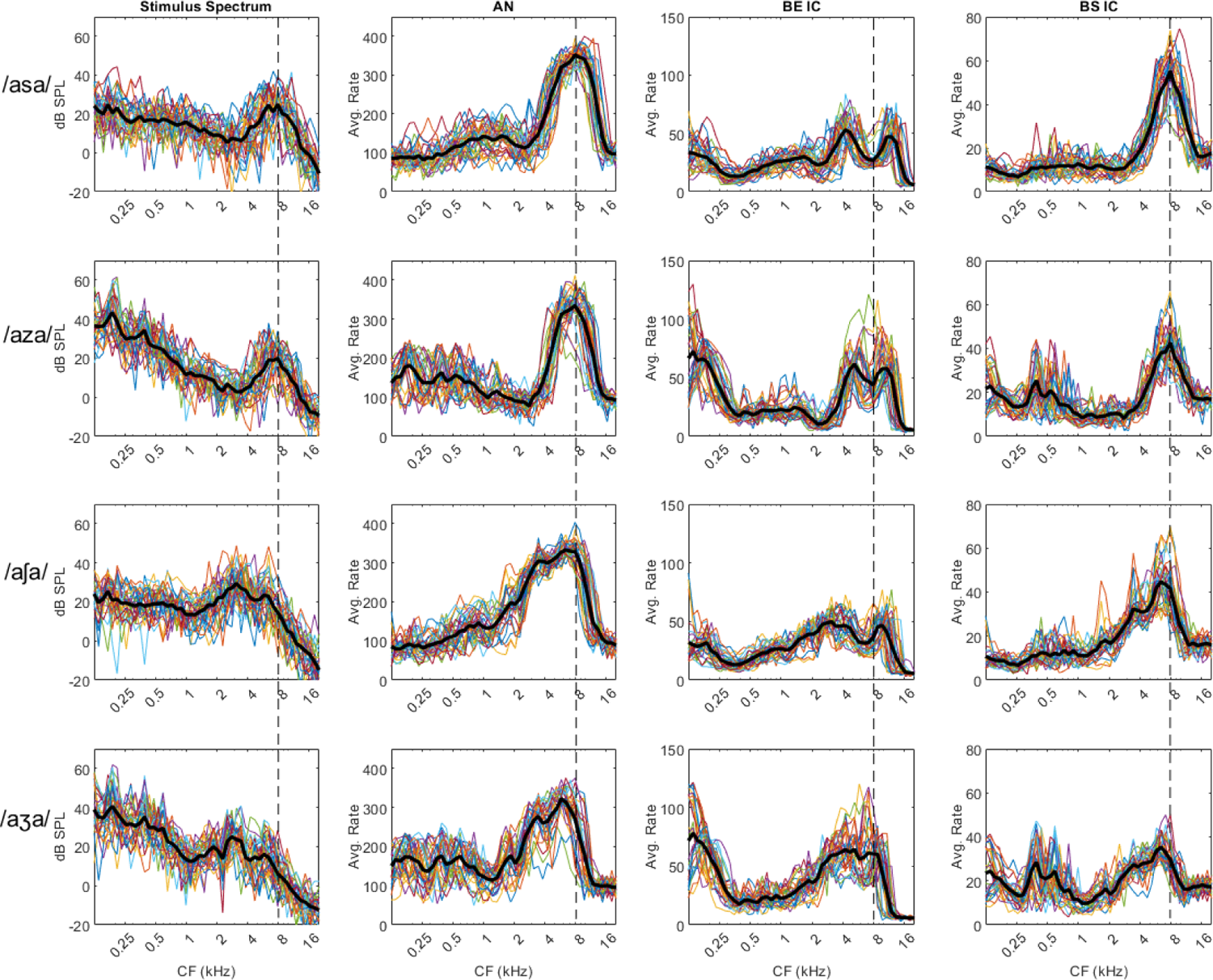
Spectral and Neural Responses to Fricative Sibilants. Mid-fricative spectrum and the corresponding neural response profiles for English sibilant fricatives. Panels from left to right demonstrate the spectra, auditory-nerve (AN), band-enhanced inferior colliculus (BE IC), and band-suppressed inferior colliculus (BS IC) responses. English sibilant fricatives from top to bottom are /s, /z/, /ʃ/, and/ʒ/. Each plot includes the 30 responses obtained from the 10 speakers. The black line indicates the average response. The vertical line at 8 kHz demarcates the boundary used for the extended high-frequency comparison.

**Figure 2.**
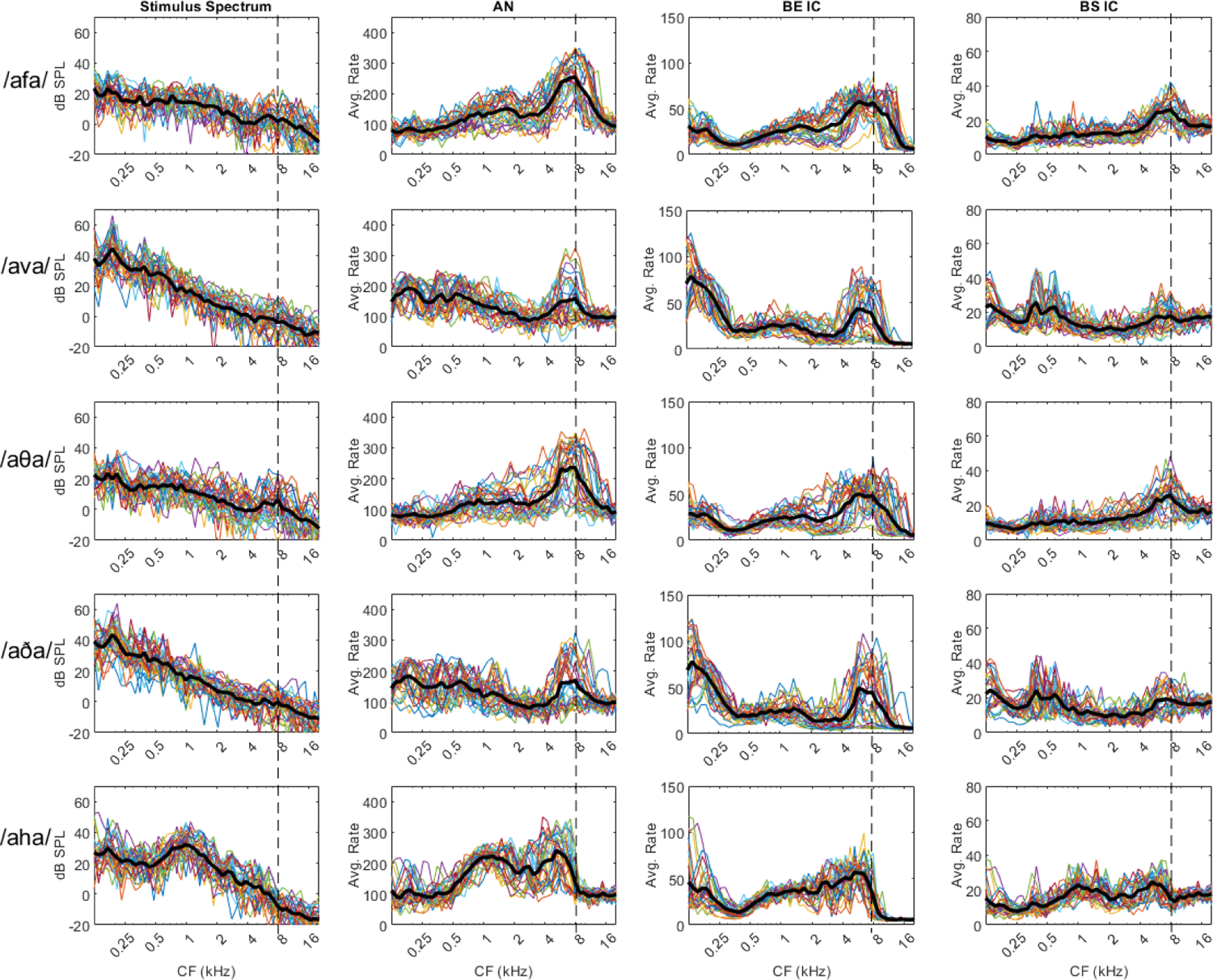
Spectral and Neural Responses to Fricative Non-Sibilants. Same format as Fig. 1 The stimuli from top to bottom are English non-sibilant fricatives /f/, /v/, /θ/, /ð/, and /h/.

**Figure 3.**
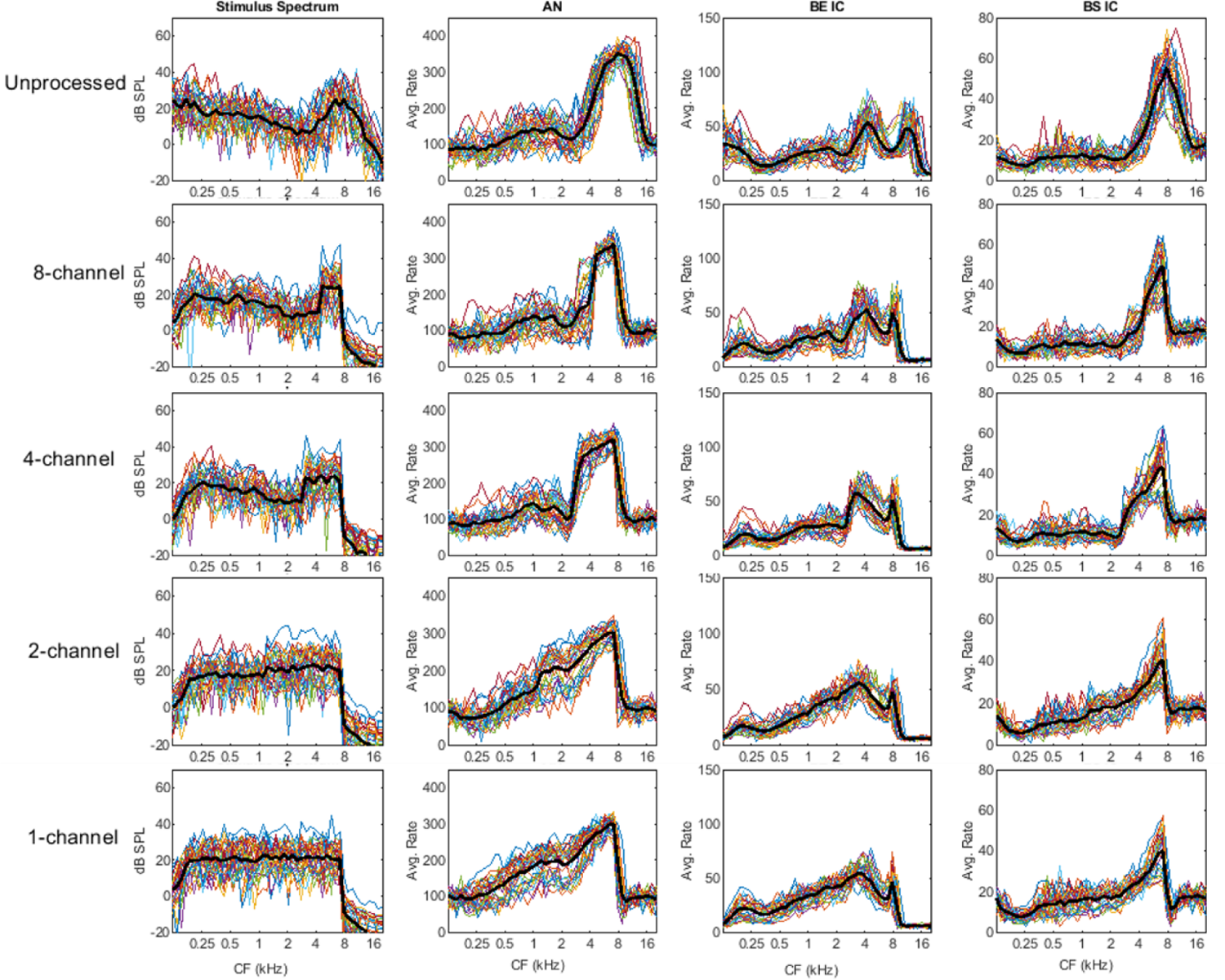
Spectral and Neural Responses to /asa/ Across the Different Conditions. Same format as Fig. 1, except that stimulus conditions increase in spectral degradation from top to bottom, starting with the unprocessed stimuli, 8-channel, 4-channel, 2-channel, and 1-channel conditions.

In the next section, the BE IC response panels will be used as a representation of the neural responses. In Figs. 1 and 2, BE IC responses encode voicing by a low-frequency peak below 250 Hz. Fricatives with the same place of articulation that differ only in voicing show similar BE IC response patterns that differ only in the low-frequency response. Sibilants show a double-peak response, while non-sibilants show a single-peak response at high frequencies. Postalveolar (/ʃ/, /ʒ/) sibilant’s first peak is broader and encompasses lower frequencies compared to alveolar sibilants (/s/, /z/). The sibilant /h/ shows a distinct absence of response beyond 8 kHz. Figure 1 shows that the unique double-peak vs. single pick response is mostly encoded by frequencies beyond 8 kHz (dashed line).

Figures 3 and 4, BE IC responses show that the processed conditions affect the voicing cue (low-frequency peak below 250 Hz), and therefore, confusions across voiced vs. voiceless fricatives with the same place of articulation are to be expected. Furthermore, the differences in the high-frequency double-peak response between the sibilants with different places of articulation (alveolar vs. postalveolar) were diminished; the first peak was broader for the processed conditions. Thus, confusions within the sibilant category would be expected to increase for the processed conditions, particularly when the number of channels was reduced. Figure 5 shows that for non-sibilant fricatives, limiting the information to approximately 8 kHz in the processed conditions would introduce confusion with the fricative /h/.

**Figure 4.**
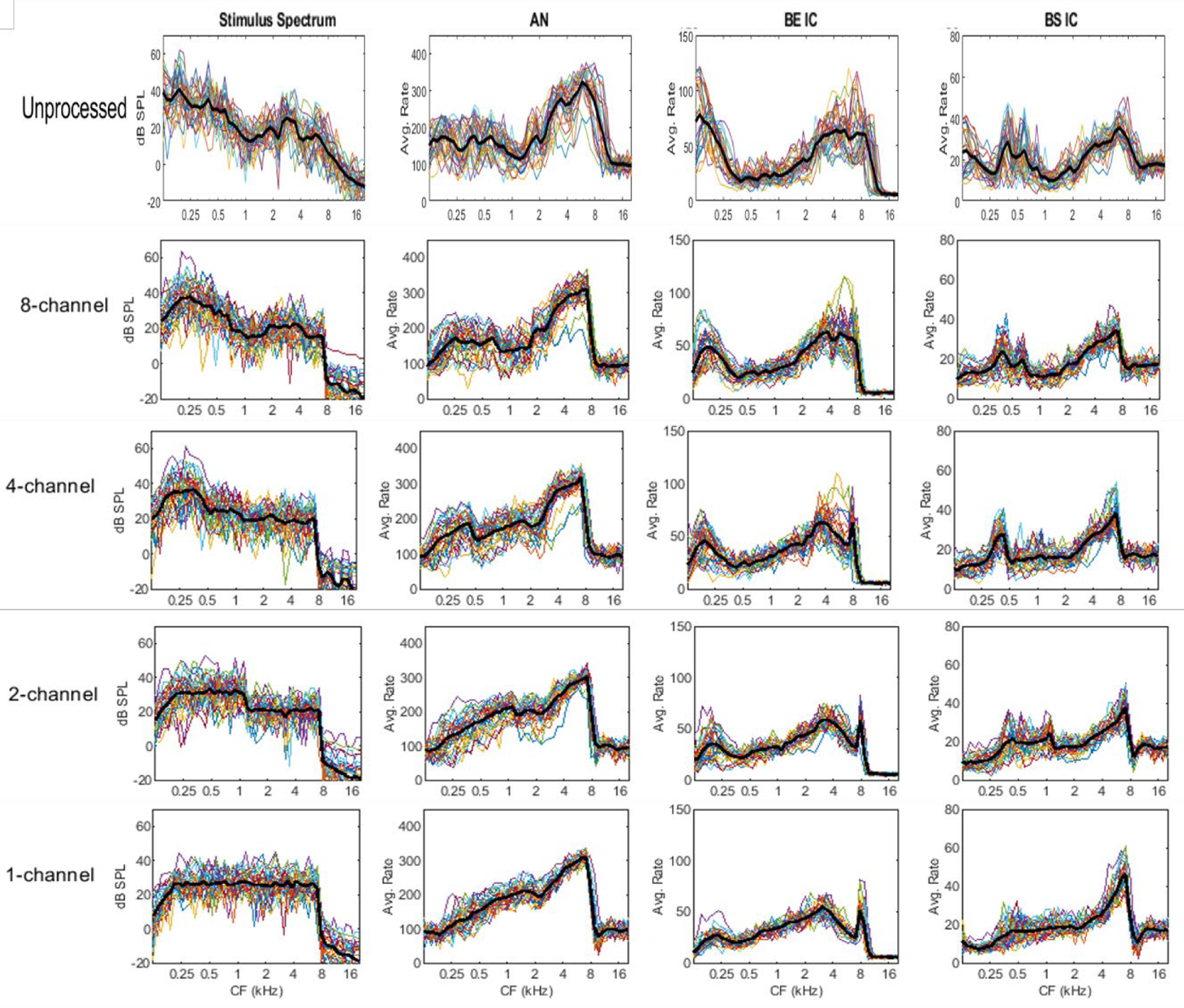
Spectral and Neural Responses to /aʒa/ Across the Different Conditions. Same format as Fig. 3.

### A. Classifier-based analysis

Tables 2I-VI demonstrate classifier-based and behavioral confusion matrices. Each table corresponds to one of the stimulus conditions. For each classifier, there were nine classes corresponding to the nine English fricatives (/f/, /v/, /θ/, /ð/, /s/, /z/, /ʃ/, /ʒ/, /h/). Tables 2I and 2II show the classifier results for the unprocessed conditions. Table 2I includes results for the extended frequency (up to 20 kHz) and Table 2II shows results for the conventional (8-kHz) frequency condition using 41 CFs. Tables 2III-VI show results for the processed conditions (with 50 CFs up to 20 kHz) using 8, 4, 2, and 1 channels, respectively. Within each table, Panels A-D correspond to spectral, AN, BE, BS, BE+BS features. Panel E demonstrates the fricative behavioral confusion matrix derived from a larger confusion matrix including 16 VCV syllables. Behavioral scores are based on published results for a group of young listeners with normal hearing (Gallun and Souza, 2008). For each classifier analysis, the mean accuracy for classification of the nine fricatives was calculated. Bold-font accuracy scores indicate that the accuracy of identifying the fricative was above the mean classifier performance. Asterisks indicate the top four confusions within a specific confusion matrix. Table 2I shows that for the unprocessed extended-frequency condition, the overall accuracy of the classifier-based analysis improved from stimulus spectrum (73.7%) to AN (83%) to BE (85.6%). On the other hand, for the unprocessed conventional-frequency and processed conditions (Tables 2II-VI), the overall accuracy of the classifier-based analysis improved from stimulus to AN model, but then decreased for the IC model results.

**Table 2I.** Classifier-based confusion matrices for the unprocessed stimuli, with 50 CFs from 125 Hz to 20 kHz.

| Classifier-Based Prediction (%) |  |  |  |  |  |  |  |  |  |
| --- | --- | --- | --- | --- | --- | --- | --- | --- | --- |
|  | /s/ | /z/ | /ʃ/ | /ʒ/ | /f/ | /v/ | /θ/ | /ð/ | /h/ |
| **stimuli** | **A- Spectral: Overall Accuracy 73.7** |  |  |  |  |  |  |  |  |
| /s/ | 80 | 3 | 3 |  | 7 |  | 7 |  |  |
| /z/ | 3 | 97 |  |  |  |  |  |  |  |
| /ʃ/ |  |  | 100 |  |  |  |  |  |  |
| /ʒ/ |  |  |  | 93 |  | 7 |  |  |  |
| /f/ | 3 |  |  |  | 63 | 3 | 30* |  |  |
| /v/ |  |  |  |  |  | 27 | 3 | 70* |  |
| /θ/ |  |  | 3 |  | 37* | 3 | 53 | 3 |  |
| /ð/ |  | 3 |  |  | 3 | 40* |  | 53 |  |
| /h/ |  |  |  |  |  |  |  | 3 | 97 |
| **B- AN: Overall Accuracy 83.0** |  |  |  |  |  |  |  |  |  |
| /s/ | 100 |  |  |  |  |  |  |  |  |
| /z/ | 3 | 97 |  |  |  |  |  |  |  |
| /ʃ/ |  |  | 100 |  |  |  |  |  |  |
| /ʒ/ |  |  |  | 97 |  | 3 |  |  |  |
| /f/ |  |  |  |  | 70 |  | 30* |  |  |
| /v/ |  | 3 |  |  |  | 53 |  | 43* |  |
| /θ/ |  |  |  |  | 17* |  | 83 |  |  |
| /ð/ |  | 3 |  |  | 3 | 47* |  | 47 |  |
| /h/ |  |  |  |  |  |  |  |  | 100 |
| **C- BE IC: Overall Accuracy 85.6** |  |  |  |  |  |  |  |  |  |
| /s/ | 100 |  |  |  |  |  |  |  |  |
| /z/ |  | 100 |  |  |  |  |  |  |  |
| /ʃ/ |  |  | 100 |  |  |  |  |  |  |
| /ʒ/ |  |  | 3 | 93 |  | 3 |  |  |  |
| /f/ |  |  |  |  | 77 |  | 23* |  |  |
| /v/ |  |  |  |  |  | 63 |  | 37* |  |
| /θ/ |  |  |  |  | 20* |  | 80 |  |  |
| /ð/ |  |  |  |  | 3 | 37* | 3 | 57 |  |
| /h/ |  |  |  |  |  |  |  |  | 100 |
| **D- BS IC: Overall Accuracy 79.6** |  |  |  |  |  |  |  |  |  |
| /s/ | 97 |  | 3 |  |  |  |  |  |  |
| /z/ | 3 | 93 |  |  |  | 3 |  |  |  |
| /ʃ/ |  |  | 100 |  |  |  |  |  |  |
| /ʒ/ |  |  |  | 97 |  |  |  | 3 |  |
| /f/ |  |  |  |  | 73 |  | 23* |  | 3 |
| /v/ |  |  |  |  | 3 | 53 |  | 43* |  |
| /θ/ |  |  | 7 |  | 37* |  | 57 |  |  |
| /ð/ |  |  |  |  | 3 | 50* |  | 47 |  |
| /h/ |  |  |  |  |  |  |  |  | 100 |
| **E- BE+ BS IC: Overall Accuracy 84.1** |  |  |  |  |  |  |  |  |  |
| /s/ | 97 |  | 3 |  |  |  |  |  |  |
| /z/ | 3 | 100 |  |  |  |  |  |  |  |
| /ʃ/ |  |  | 100 |  |  |  |  |  |  |
| /ʒ/ |  |  |  | 97 |  | 3 |  |  |  |
| /f/ |  |  |  |  | 80 |  | 20* |  |  |
| /v/ |  |  |  |  |  | 70 |  | 30* |  |
| /θ/ |  |  | 3 |  | 33* |  | 63 |  |  |
| /ð/ |  | 3 |  |  | 3 | 40* |  | 53 |  |
| /h/ |  |  |  |  | 3 |  |  |  | 97 |
| **F-Behavioral: Overall Accuracy 90.8** |  |  |  |  |  |  |  |  |  |
| /s/ | 99 |  |  |  |  |  |  | 1 |  |
| /z/ |  | 100 |  |  |  |  |  |  |  |
| /ʃ/ |  |  | 100 |  |  |  |  |  |  |
| /ʒ/ |  |  |  | 100 |  |  |  |  |  |
| /f/ | 6 |  | 15* |  | 79 |  |  |  |  |
| /v/ |  |  |  |  |  | 98 |  | 2 |  |
| /θ/ | 19* |  |  |  | 8* |  | 66 | 6 | 1 |
| /ð/ |  | 3 |  |  |  | 5 | 7* | 85 |  |

**Table 2II.**
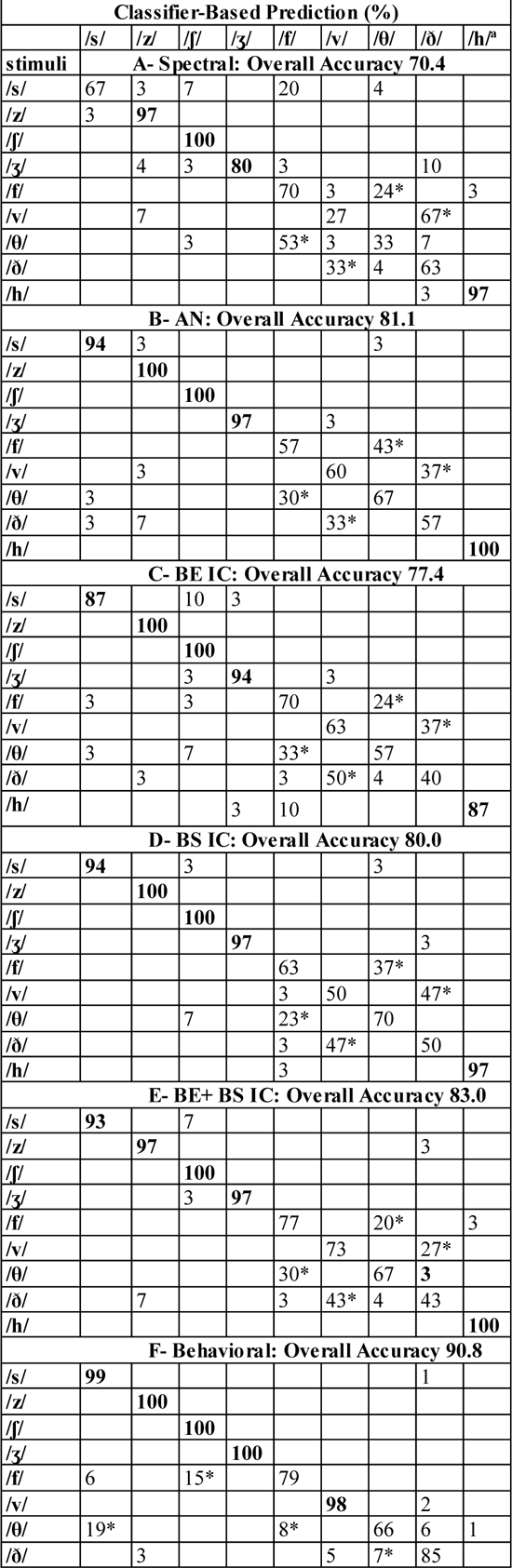
Classifier-based confusion matrices using the unprocessed stimuli with 41 CFs from 125 Hz to 8 kHz.

**Table 2III.** Classifier-based confusion matrices using the 8-channel processed stimuli, with 50 CFs from 125 Hz to 20 kHz.

| Classifier-Based Prediction (%) |  |  |  |  |  |  |  |  |  |
| --- | --- | --- | --- | --- | --- | --- | --- | --- | --- |
|  | /s/ | /z/ | /ʃ/ | /ʒ/ | /f/ | /v/ | /θ/ | /ð/ | /h/ |
| stimuli | A- Spectral: Overall Accuracy 50.3 |  |  |  |  |  |  |  |  |
| /s/ | 30 |  | 33* | 3 | 20 |  |  |  | 13 |
| /z/ | 20 | 33 |  | 3 |  | 20 | 3 | 13 | 7 |
| /ʃ/ |  |  | 100 |  |  |  |  |  |  |
| /ʒ/ |  | 3 | 17 | 73 |  |  |  | 3 | 3 |
| /f/ |  |  | 7 |  | 23 |  | 3 |  | 67* |
| /v/ |  |  |  |  |  | 40 | 27 | 30 | 3 |
| /θ/ |  |  | 3 |  | 33* | 3 | 7 |  | 53* |
| /ð/ |  |  |  |  | 7 | 20 | 20 | 50 | 3 |
| /h/ |  |  | 3 |  |  |  |  |  | 97 |
| B- AN: Overall Accuracy 59.2 |  |  |  |  |  |  |  |  |  |
| /s/ | 7 |  | 17 |  | 53* |  | 20 |  | 3 |
| /z/ | 3 | 73 | 3 | 3 |  | 10 | 3 | 3 |  |
| /ʃ/ |  |  | 100 |  |  |  |  |  |  |
| /ʒ/ |  |  | 23 | 70 | 3 |  |  |  | 3 |
| /f/ |  |  |  |  | 30 |  | 47* |  | 23 |
| /v/ |  |  |  |  | 0 | 60 | 33* | 7 |  |
| /θ/ |  |  | 7 |  | 17 |  | 63 |  | 13 |
| /ð/ |  |  |  |  |  | 37* | 33* | 30 |  |
| /h/ |  |  |  |  |  |  |  |  | 100 |
| C- BE IC: Overall Accuracy 37.8 |  |  |  |  |  |  |  |  |  |
| /s/ | 7 |  | 53* |  | 3 |  | 13 |  | 23 |
| /z/ | 3 | 7 | 7 |  | 7 | 33* | 27 | 7 | 10 |
| /ʃ/ |  |  | 90 |  |  |  |  |  | 10 |
| /ʒ/ |  |  | 13 | 20 |  |  |  |  | 67* |
| /f/ |  |  | 3 |  | 3 |  | 33* |  | 60* |
| /v/ |  |  |  |  |  | 23 | 23 | 33* | 20 |
| /θ/ |  |  | 10 |  |  |  | 57 |  | 33 |
| /ð/ |  |  |  |  |  | 20 | 27 | 33 | 17 |
| /h/ |  |  |  |  |  |  |  |  | 100 |
| D- BS IC: Overall Accuracy 55.2 |  |  |  |  |  |  |  |  |  |
| /s/ | 20 |  | 47* |  | 23 |  | 10 |  |  |
| /z/ | 3 | 40 | 13 |  | 10 | 7 | 7 | 20 |  |
| /ʃ/ |  |  | 100 |  |  |  |  |  |  |
| /ʒ/ |  | 3 | 33* | 57 |  | 3 |  |  | 3 |
| /f/ |  |  |  |  | 73 |  | 20 |  | 7 |
| /v/ |  |  |  |  | 17 | 37 | 27 | 20 |  |
| /θ/ |  |  | 3 |  | 43* |  | 43 |  | 10 |
| /ð/ |  |  |  |  | 20 | 17 | 33* | 30 |  |
| /h/ |  |  |  |  | 3 |  |  |  | 97 |
| E- BE+ BS IC: Overall Accuracy 41.4 |  |  |  |  |  |  |  |  |  |
| /s/ | 0 |  | 70* |  | 3 |  | 13 |  | 13 |
| /z/ | 7 | 10 | 7 |  | 10 | 33* | 20 | 3 | 10 |
| /ʃ/ |  |  | 100 |  |  |  |  |  |  |
| /ʒ/ |  |  | 20 | 40 |  |  |  |  | 40* |
| /f/ |  |  | 7 |  | 0 |  | 30 |  | 63* |
| /v/ |  |  |  |  |  | 30 | 20 | 33* | 17 |
| /θ/ |  |  | 13 |  |  |  | 53 |  | 33* |
| /ð/ |  |  |  |  |  | 23 | 23 | 40 | 13 |
| /h/ |  |  |  |  |  |  |  |  | 100 |
| F- Behavioral: Overall Accuracy 69.4 |  |  |  |  |  |  |  |  |  |
| /s/ | 81 | 3 | 3 |  | 7 |  | 6 |  |  |
| /z/ | 3 | 70 |  | 9 |  | 6 |  | 12 |  |
| /ʃ/ | 1 |  | 99 |  |  |  |  |  |  |
| /ʒ/ |  |  |  | 100 |  |  |  |  |  |
| /f/ | 10 |  |  |  | 69 | 1 | 20* |  |  |
| /v/ |  | 6 |  |  |  | 59 |  | 23* | 12 |
| /θ/ | 19 |  | 1 |  | 46* |  | 31 | 1 | 2 |
| /ð/ |  | 6 |  |  |  | 27* | 6 | 46 | 15 |

**Table 2IV.** Classifier-based confusion matrices using the 4-channel processed stimuli, with 50 CFs from 125 Hz to 20 kHz.

| Classifier-Based Prediction (%) |  |  |  |  |  |  |  |  |  |
| --- | --- | --- | --- | --- | --- | --- | --- | --- | --- |
|  | /s/ | /z/ | /ʃ/ | /ʒ/ | /f/ | /v/ | /θ/ | /ð/ | /h/ |
| **stimuli** | **A- Spectral: Overall Accuracy 36.1** |  |  |  |  |  |  |  |  |
| /s/ | 3 |  | 77* |  | 3 |  |  |  | 17 |
| /z/ | 10 | 3 | 17 | 47* |  | 3 |  | 10 | 10 |
| /ʃ/ |  |  | **93** |  |  |  |  |  | 7 |
| /ʒ/ |  |  | 27 | **37** |  | 3 |  | 3 | 30 |
| /f/ |  |  |  |  | 23 |  |  |  | 77* |
| /v/ |  |  |  | 7 |  | 30 | 17 | 20 | 27 |
| /θ/ |  |  | 7 | 3 | 37 |  | 13 |  | 40* |
| /ð/ |  |  |  | 7 | 7 | 27 | 20 | 23 | 17 |
| /h/ |  |  |  |  |  |  |  |  | **100** |
| **B- AN: Overall Accuracy 47.3** |  |  |  |  |  |  |  |  |  |
| /s/ | 0 |  | 57* |  | 30 |  | 10 |  | 3 |
| /z/ |  | 10 | 13 | 50* | 3 |  |  | 17 | 7 |
| /ʃ/ |  |  | **100** |  |  |  |  |  |  |
| /ʒ/ |  |  | 33 | 43* | 3 |  |  |  | 20 |
| /f/ |  |  |  |  | 37 |  | 47* |  | 17 |
| /v/ |  |  |  | 3 | 3 | **63** | 23 | 7 |  |
| /θ/ |  |  | 7 |  | 30 |  | **60** |  | 3 |
| /ð/ |  |  |  | 3 | 7 | 40 | 37 | 13 |  |
| /h/ |  |  |  |  |  |  |  |  | **100** |
| **C- BE IC: Overall Accuracy 29.2** |  |  |  |  |  |  |  |  |  |
| /s/ | 0 |  | 60* |  | 3 |  | 7 |  | 30 |
| /z/ |  | 0 | 33 | 3 |  |  |  |  | 63* |
| /ʃ/ |  |  | **100** |  |  |  |  |  |  |
| /ʒ/ |  |  | 27 | 7 |  |  |  |  | 67* |
| /f/ |  |  | 3 |  | 3 |  | 30 |  | 63* |
| /v/ |  |  |  |  |  | 0 | 27 | 27 | 47 |
| /θ/ |  |  | 3 |  |  |  | **57** |  | 40 |
| /ð/ |  |  |  |  |  | 3 | 33 | 13 | 50 |
| /h/ |  |  | 17 |  |  |  |  |  | **83** |
| **D- BS IC: Overall Accuracy 32.6** |  |  |  |  |  |  |  |  |  |
| /s/ | 0 |  | 67* |  | 27 |  | 7 |  |  |
| /z/ |  | 0 | 37 | 23 | 13 |  | 3 | 20 | 3 |
| /ʃ/ |  |  | **100** |  |  |  |  |  |  |
| /ʒ/ |  |  | 37 | **43** | 7 |  |  |  | 13 |
| /f/ |  |  |  |  | **70** |  | 27 |  | 3 |
| /v/ |  |  |  |  | 17 | **33** | 2 | 27 | 3 |
| /θ/ |  |  | 3 |  | 43* |  | **47** |  | 7 |
| /ð/ |  |  |  |  | 20 |  |  | 0 | 80* |
| /h/ |  |  |  |  | 17 | 27 | 17 | 40* | 0 |
| **E- BE+ BS IC: Overall Accuracy 34.1** |  |  |  |  |  |  |  |  |  |
| /s/ | 10 |  | 70* |  |  |  | 7 |  | 13 |
| /z/ |  | 0 | 33 | 13 |  |  | 3 |  | 50* |
| /ʃ/ |  |  | **100** |  |  |  |  |  |  |
| /ʒ/ |  |  | 40 | 20 |  |  |  |  | 40 |
| /f/ |  |  | 3 |  | 0 |  | 30 |  | 67* |
| /v/ |  |  |  |  |  | 20 | 23 | 17 | 40 |
| /θ/ |  |  | 17 |  |  |  | **50** |  | 33 |
| /ð/ |  |  |  |  |  | 10 | 37 | 10 | 43* |
| /h/ |  |  | 3 |  |  |  |  |  | **97** |
| **F- Behavioral: Overall Accuracy 46.6** |  |  |  |  |  |  |  |  |  |
| /s/ | **53** | 2 | 19 |  | 13 | 2 | 11 |  |  |
| /z/ | 4 | **54** |  | 22 |  | 15 | 1 | 4 |  |
| /ʃ/ | 49* |  | **49** |  | 2 |  |  |  |  |
| /ʒ/ | 4 | 33* | 2 | **54** |  | 5 |  | 2 |  |
| /f/ | 9 |  |  |  | **63** |  | 22 | 4 | 2 |
| /v/ | 1 | 2 |  |  |  | **64** |  | 9 | 24 |
| /θ/ | 14 | 1 | 2 |  | 49* | 5 | 24 | 2 | 3 |
| /ð/ |  | 2 |  |  | 1 | 65* | 3 | 14 | 15 |

**Table 2V.**
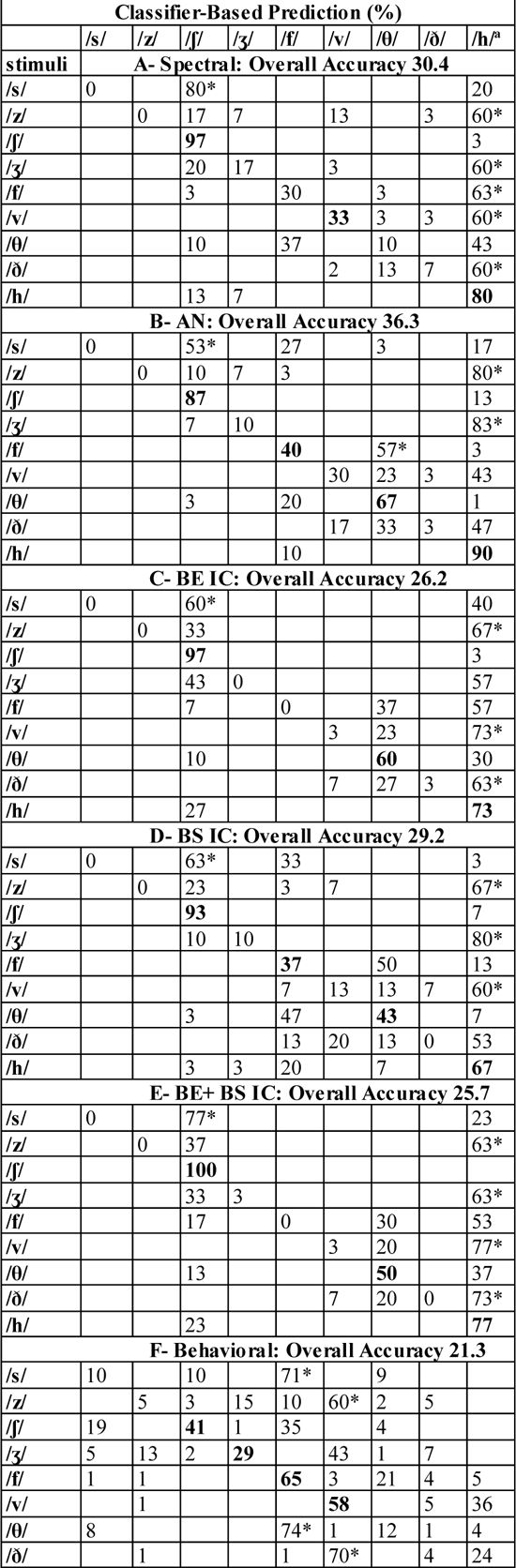
Classifier-based confusion matrices using the 2-channel processed stimuli, with 50 CFs from 125 Hz to 20 kHz.

**Table 2VI.**
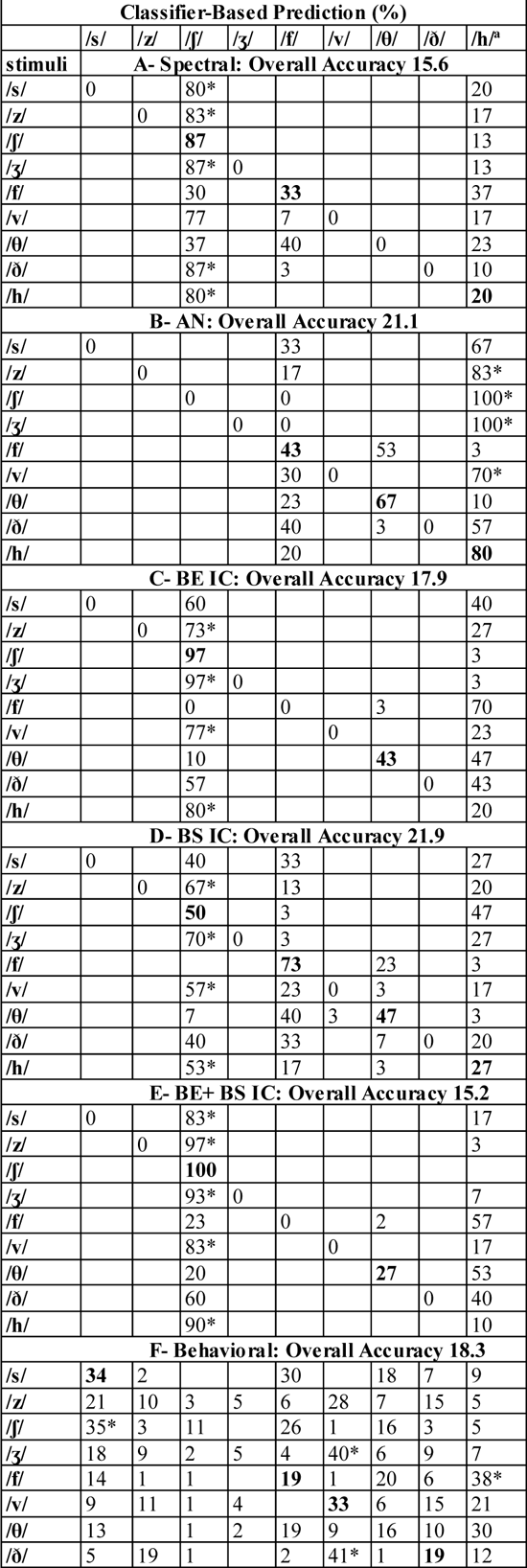
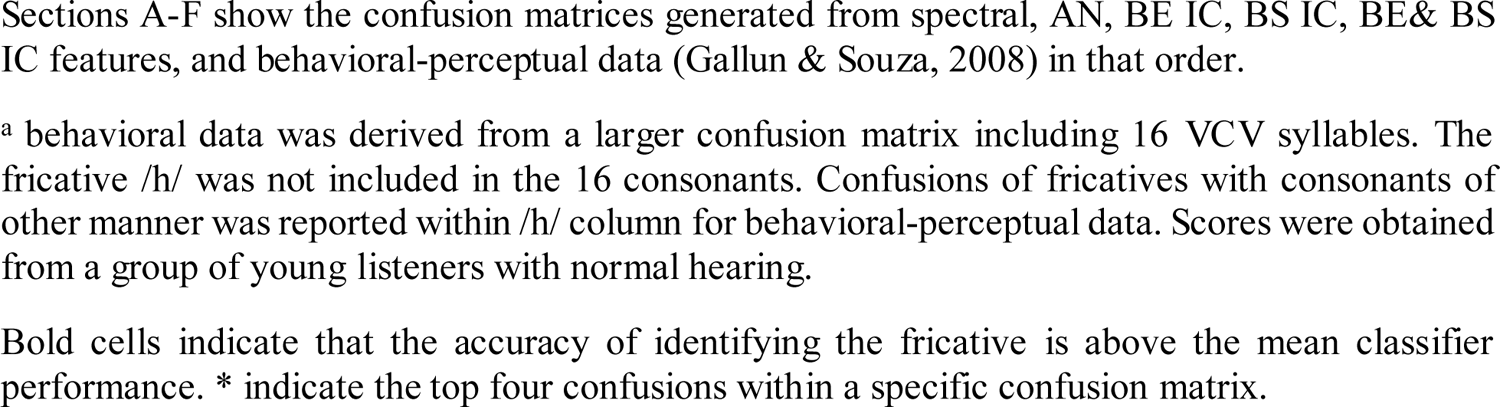
Classifier-based confusion matrices using the 1-channel processed stimuli, with 50 CFs from 125 Hz to 20 kHz.

Compared to extended frequency data (Table 2I), limiting the data to 8 kHz reduced the overall accuracy for the spectral (73.7% vs. 70.4%), AN (83.0% vs. 81.1%),BE (85.6% vs. 77.4%) and BE+ BS (84.1% vs. 83.0%) models, but not the BS IC model (79.6% vs. 80.4%). The BE IC model showed the largest effect. For the neural response models, apart from the differences in the overall accuracy between the 20 and 8 kHz models, no noticeable differences were seen in the patterns of accuracies and confusions, i.e., in the 8-kHz model, fricatives were less accurately identified, but the pattern of accuracies was the same as for the 20-kHz model. For the spectral analysis, limiting the information to 8 kHz resulted in more confusions of the sibilant /s/.

In the unprocessed condition, the highest accuracy was achieved by the BE features, whereas in the processed conditions, the BS or BE+BS features were more accurate than BE. Classifier based accuracies approached behavioral accuracy (90%) for the unprocessed, and 4- and 8-channel processed conditions. For the 2- and 1-channel processed conditions, model-based accuracies were slightly higher than the behavioral accuracies.

Looking closely at the fricative-specific accuracy scores in the unprocessed condition, modeled neural responses provided robust correlates to behavioral accuracy. Sibilants were predicted with higher accuracies relative to non-sibilants (apart from /h/), similar to the behavioral accuracies, with the exception of /v/. However, in terms of fricative-specific confusions, in the modeled neural responses it was rare to confuse a non-sibilant with a sibilant, unlike the behavioral data. Both the modeled neural responses and behavioral data showed the highest confusions among the non-sibilant fricatives; however, in the modeled neural responses, confusions were limited to within the non-sibilant fricatives (/f/ with /θ/ and /v/ with /ð/), whereas for the behavioral data, confusions of non-sibilant fricatives with sibilant fricatives were most common (/θ/ with /s/ and /f/ with /ʃ/). The voicing cue was robust in the classifier-based analysis, with less confusions between voiced and voiceless fricatives. However, in the behavioral data, confusion of /ð/ with /θ/, a voicing contrast, was among the four most common confusions.

In the processed conditions, sibilant accuracy appeared to be more affected than non-sibilant, as predicted from Fig. 3, where the differences in the high-frequency double-peak response between the sibilants with different places of articulation (alveolar vs. postalveolar) were diminished. This decline in sibilant accuracy increased for the degraded conditions, however, this trend was not in line with the behavioral accuracy for the less-degraded conditions (8 and 4-channels), where sibilants were more accurately identified. The discrepancy between the modeled and behavioral results in less-degraded conditions may highlight the importance of the higher order processing in the difficult listening conditions and that those capacities have a limit (less vs. more degraded). For confusions, as anticipated from Fig. 5 for non-sibilant fricatives, limiting the information to 8 kHz, to match the frequency range used in the behavioral task, resulted in confusions with the fricative /h/. The behavioral task lacked the fricative /h/.

**Figure 5.**
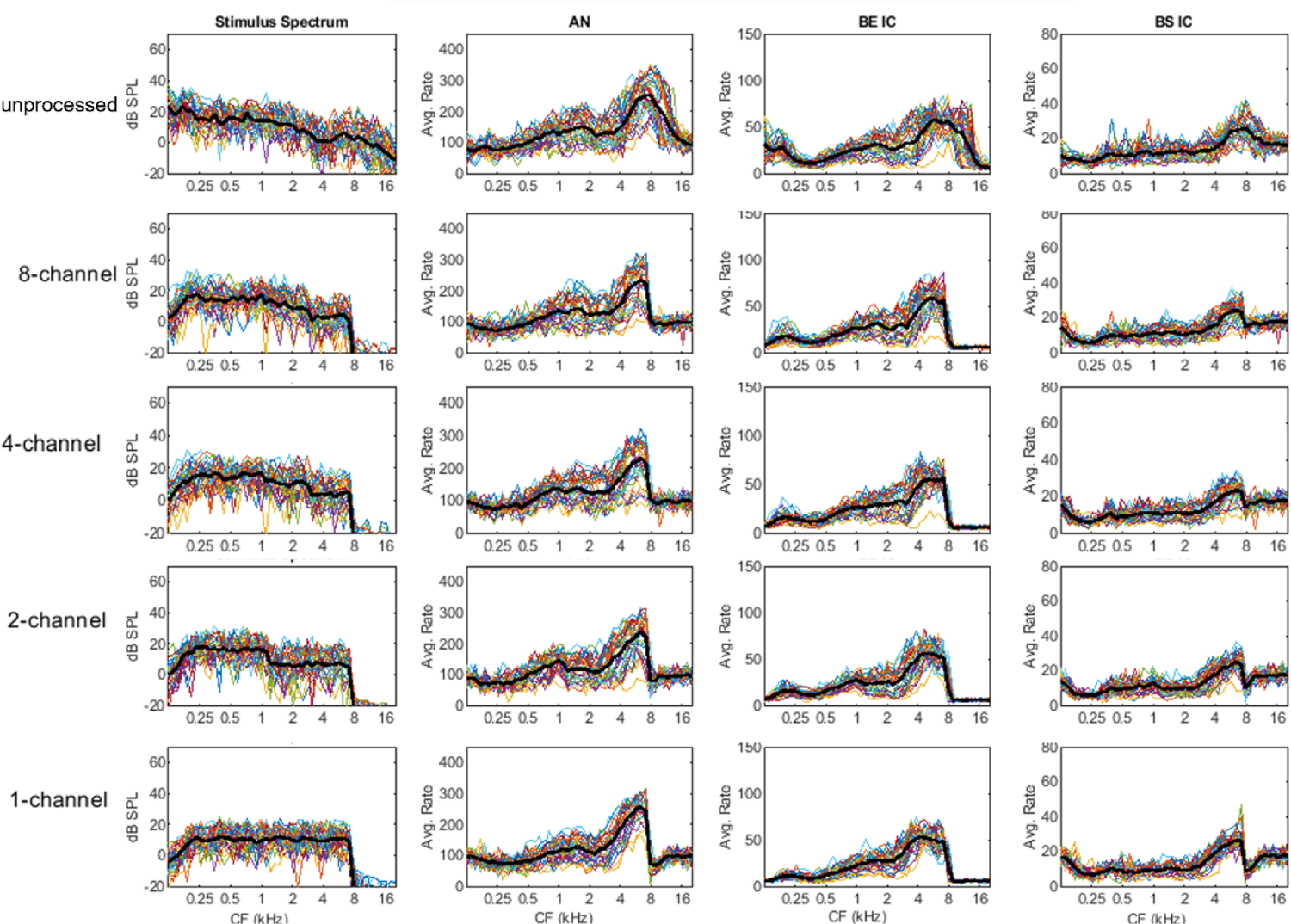
Spectral and Neural Responses to /afa/ Across the Different Conditions. Same format as Fig. 3.

## IV. DISCUSSION

Fricatives have been shown to exhibit considerable contextual and intra/inter-subject variability (Jongman et al., 2000; Narayanan et al., 1995; Shadle, 1990), as well as asymmetries in fricative voiced and voiceless contrast patterns cross linguistically (Chodroff and Wilson, 2020; Maddieson, 1984; Stevens, 1998). Yet, despite this variability and these patterns, fricative contrasts are perceived with high accuracy (Cutler et al., 2004; Gallun and Souza, 2008; Pisoni and Luce, 1986; Woods et al., 2010). To understand how, in the face of the variability in production, consonant perception is robust, research has focused on the production system, the articulation and acoustic analysis of fricatives. The produced waveform is a mechanical signal transformed at the cochlea and AN into a neural signal that moves through the complex neural pathways of the auditory system. While the acoustic analyses are a powerful and productive tool, they lend little insight into the neural coding of speech, or the constraints imposed on speech perception by the auditory system. Realistic neural models have been developed to model the coding of speech in the midbrain and may help in better understanding the link between acoustical signals and behavioral accuracy (Carney, 2018; Carney and McDonough, 2019; Nelson and Carney, 2004; Zilany et al., 2014; Zilany et al., 2009).

The current study used computational models for AN fibers and IC neurons to demonstrate the auditory representation of fricatives and establish the link between the production and perception of fricatives. The modeled neural responses, from 10 different speakers uttering each fricative 3 times, were compared to behavioral accuracy from a study by Gallun and Souza (2008). We hypothesized that the modeled neural responses can aid in developing explanations and representations of behavioral accuracies than the spectra of the acoustical signal alone.

### A. Classifier vs. perceptual performance

The classifier analysis supported our hypothesis in the unprocessed condition (natural speech), for which the overall accuracy improved from the stimulus (73.7%) to AN (83%) to IC (BE, 85.6%), for the 20-kHz bandwidth conditions, approaching the behavioral accuracy (90%). The neural modeled accuracies remained below the behavioral accuracies, as expected assuming the behavioral task benefits from higher-level neural equivalents of categorical perception (Lago et al., 2015), and top-down processing (Davis and Johnsrude, 2007). If anything, a larger gap between the classifier accuracy and behavioral accuracy was expected. The high accuracy in the classifier results could be explained by the smaller number of phoneme choices in the classifier (n=9) vs. perceptual (n=16) task.

For the processed conditions, the overall accuracy of the classifier analysis improved from the stimulus spectra to AN but then declined at the level of the IC, which could imply a greater role of top-down processing in degraded conditions. Top-down processing is thought to involve selective favoring of features based on prior knowledge. Highly-weighted features influence the processing and encoding of sensory input (Asilador and Llano, 2021; Von Helmholtz, 1867). The neural modeled accuracies remained below the behavioral accuracies for less degraded conditions (8- and 4-channel). However, for the more degraded conditions (2- and 1-channel), behavioral accuracies were slightly lower than the neural modeled accuracies. The higher accuracy in the classifier-based models in the more degraded conditions could be, once again, explained by the difference in task design (larger set in the perceptual task) and by the limits of higher order processing for strongly degraded stimuli.

### B. Fricative-specific performance

The fricative-specific accuracy scores of the modeled neural responses were consistent with the behavioral accuracy. Sibilants were predicted more accurately than non-sibilants. Classifier-based accuracy scores matched the behavioral data (apart from /v/). Other behavioral studies have shown controversial results as to whether /v/ is highly confusable (Cutler et al., 2004; Woods et al., 2010). The unprocessed classifier confusions were not consistent with the behavioral data, suggesting that different processes may be involved in accuracy vs. confusions. While the accuracy of identification of a fricative may rely on encoding at the level of the IC, the pattern of fricative confusions might rely on other processes, including language phonotactics, experience, and top-down processing. Meyer et al. (2010) has shown that human phoneme recognition depending on speech-intrinsic variability was associated with different predictors for recognition rates vs. phoneme confusions.

The results for different IC cell types in the classifier analysis varied with the acoustic signal quality; thus, the relative role of different IC cell types may vary in different listening environments. For the unprocessed condition, the BE features yielded the highest accuracy, whereas, in the processed conditions, the BS or BE+BS IC features yielded the highest accuracy. The two groups of IC neurons represent stimulus features with opposite polarities, similar to the retinal ganglion cells with on-center-off-surround and off-center-on-surround receptive fields (Carney and McDonough, 2019; Kim et al., 2020).

### C. Extended high-frequency hearing

In the current study, we compared the classifier performance for the spectra or model responses limited to 8 kHz and 20 kHz to understand the role of extended frequencies on accuracy at different levels from stimulus to the IC. Results showed that extended frequencies primarily affected the accuracy performance based on the BE IC model neurons. It is generally assumed that frequencies limited to 8 kHz contain much of the phonetic cues, which has reduced the focus on the importance of the extended frequencies (Vitela et al., 2015). The results based on the stimulus spectra and AN models benefited minimally from the additional information provided by the extended frequencies, whereas the BE IC results did. Limiting the frequency information did not seem to affect the fricative-specific accuracies and confusions, i.e., it reduced the overall accuracy equally for the neural responses but affected the accuracies of the sibilant /s/ based on spectral infomration. The current study corroborated the growing evidence suggesting that acoustic information in the higher-frequency regions affects speech intelligibility and that the association between the spectral information and neural responses is not straightforward (Badri et al., 2011; Levy et al., 2015; Moore et al., 2017; Pienkowski, 2017; Zadeh et al., 2019).

### D. Conclusions, limitations, and future work

The current study showed that the modeled neural responses at the level of AN and IC provided better predictions of behavioral accuracies than did the stimulus spectra. Accuracies of fricative contrasts were explained by modeled neural response profiles, while confusions were only partially explained. Different IC cell types may play different roles in consonant perception, based on the quality of the acoustic signals. Future studies measuring electrophysiological responses from different IC cell types may shed light on these roles. Extended frequencies (8-20 kHz) improved accuracies primarily for the model BE IC neurons, potentially explaining some of the discrepancy between acoustical and speech perception data for listeners with hearing loss at extended frequencies.

It is important to note that the modeled neural responses were limited to a single sound level. It is possible that by varying the sound level over a wide (realistic) range, the models could perform better, in comparison to the AN. However, behavioral data for comparison to model responses at different sound levels were not available. Other model limitations include the strictly on-CF implementation of the SFIE model and the lack of efferent control of cochlear gain, which might impact performance for degraded conditions (Farhadi et al., 2021). Comparison of the confusions in the processed conditions was limited by the frequency limit of 7.2 kHz used in the behavioral task, which resulted in higher confusions for the fricative /h/ (the behavioral data lacked the fricative /h/). Future studies that simulate neural responses of the hearing-impaired ear to fricatives could guide the development of processing strategies aimed at reaching the normal-hearing neural targets.

## Acknowledgements

Supported by NIH-DC010813 (LHC).

